# ThiL is a valid antibacterial target that is essential for both thiamine biosynthesis and salvage pathway in *Pseudomonas aeruginosa*

**DOI:** 10.1101/2020.03.04.976639

**Authors:** Hyung Jun Kim, Hyunjung Lee, Yunmi Lee, Inhee Choi, Yoonae Ko, Sangchul Lee, Soojin Jang

## Abstract

Thiamine pyrophosphate (TPP) is an essential cofactor for various pivotal cellular processes in all living organisms, including bacteria. As thiamine biosynthesis occurs in bacteria but not humans, bacterial thiamine biosynthesis is an attractive target for antibiotic development. Among enzymes in the thiamine biosynthetic pathway, thiamine monophosphate kinase (ThiL) catalyzes the final step of the pathway, phosphorylating thiamine monophosphate (TMP) to produce TPP. In this work, we extensively investigated ThiL in *Pseudomonas aeruginosa*, a major pathogen of hospital-acquired infections. We demonstrated that *thiL* deletion abolishes not only thiamine biosynthesis but also thiamine salvage capability, showing growth defects of the Δ*thiL* mutant even in the presence of thiamine derivatives except TPP. Most importantly, the pathogenesis of the Δ*thiL* mutant was markedly attenuated compared to wild-type bacteria, with lower inflammatory cytokine induction and 10^3^~10^4^ times decreased bacterial load in an *in vivo* infection model where the intracellular TPP level is in the submicromolar range. In order to validate *P. aeruginosa* ThiL (PaThiL) as a new drug target, we further characterized its biochemical properties determining a Vmax of 4.0±0.2 nomol·min^−1^ and K_M_ values of 111±8 and 8.0±3.5μM for ATP and TMP, respectively. A subsequent *in vitro* small molecule screening identified PaThiL inhibitors including WAY213613 that is a noncompetitive inhibitor with a Ki value of 13.4±2.3 μM and a potential antibacterial activity against *P. aeruginosa*. This study proved that PaThiL is a new drug target against *P. aeruginosa* providing comprehensive biological and biochemical data that could facilitate to develop a new repertoire of antibiotics.

---

*Pseudomonas aeruginosa* is a gram-negative opportunistic pathogen associated with a wide range of acute and chronic infections of various body sites, including the urinary tract, skin, and respiratory tract (1). Patients with compromised immune defenses due to underlying diseases such as cancer or HIV infection or with severe burns, cystic fibrosis, bronchiectasis, or chronic obstructive pulmonary disease are particularly susceptible to *P. aeruginosa* infection (2–7). The mainstay of treatment for *P. aeruginosa* infection is antibiotics. However, the expression of multiple efflux pumps, reduced permeability of the outer membrane, capacity to form biofilm, and the presence of persisters have rendered *P. aeruginosa* intrinsically resistant to many antibiotics (8,9). Given this natural resistance, excessive use of antibiotics is required to treat *P. aeruginosa* infections, accelerating the development of drug-resistant strains. Indeed, multidrug-resistant and extensively drug-resistant *P. aeruginosa* strains are now prevalent worldwide, with the emergence of pan-drug resistant strains threatening global public health (10–13). The World Health Organization recently listed drug-resistant *P. aeruginosa* as a priority pathogen necessitating urgent action to develop novel antibiotics to overcome current antibiotic resistance (14).

Thiamine is a crucial molecule in all living organisms, from microorganisms to mammals. Therefore, thiamine metabolism has attracted growing attention in the development of various drugs, including antibiotics (15–19). The physiologically active form of thiamine, thiamine pyrophosphate (TPP), plays important roles as a cofactor in various essential cellular processes, including carbohydrate, lipid, and amino acid metabolism (20–22). Microorganisms such as bacteria and fungi as well as plants produce TPP via de novo biosynthetic pathways that mammals lack (23,24). The TPP biosynthetic pathway of bacteria involves the separate biosynthesis of thiazole and pyrimidine moieties that are joined to form thiamine monophosphate (TMP) in a reaction catalyzed by thiamine phosphate synthase (ThiE) (25–27). Thiamine monophosphate kinase (ThiL) catalyzes the final step of the pathway by phosphorylating TMP to TPP, the biologically active form of the cofactor (28,29). In addition to the TPP biosynthetic pathway, bacteria are capable of salvaging thiamine from exogenous sources to generate TPP. Some bacteria, such as *Bacillus subtilis*, take up exogenous thiamine and convert it to TPP in a one-step reaction catalyzed by thiamine pyrophosphate kinase (TPK), similar to the mammalian thiamine salvage pathway (30–33). Other bacteria, including *Escherichia coli*, first convert thiamine to TMP by thiamine kinase (ThiK) and then subsequently generate TPP by adding one additional phosphate to TMP through ThiL, the enzyme in the main TPP biosynthetic pathway (30,34). Because TPP is indispensable for bacterial survival and humans lack the TPP biosynthetic pathway, enzymes involved in bacterial TPP biosynthesis are potential targets in the development of new antibiotics. ThiL is of particular interest considering its key role in thiamine metabolism. Nevertheless, few studies involving a limited number of bacterial species have been conducted on ThiL, and it has never been validated as a target for antibacterial agents (28,29,35,36).

In this work, we constructed a clean *thiL* deletion mutant of *P. aeruginosa* and investigated the impact of *thiL* deficiency on bacterial survival and *in vivo* pathogenesis. We also biochemically characterized *P. aeruginosa* ThiL and identified small molecules that inhibit the enzyme using an optimized luminescent kinase assay. To the best of our knowledge, this is the first work to demonstrate the role of *P. aeruginosa* ThiL in bacterial physiology and pathogenesis and the first to validate ThiL as a new target for drug development, providing comprehensive biochemical characterization of ThiL and identifying its inhibitors.

## Results

### The role of ThiL in P. aeruginosa thiamine metabolism

To investigate the physiological role of ThiL in *P. aeruginosa*, we constructed an in-frame deletion mutant of the *thiL* gene in *P. aeruginosa* PAO1 by two-step allele exchange (37–39). TPP was supplied via selection media for the last step of mutant generation to avoid the loss of bacterial viability due to impaired thiamine biosynthesis. Deletion of *thiL* in the bacterial genome was confirmed by amplification of the *thiL* flanking region (Fig. S1). Phenotypic analysis of the Δ*thiL* mutant confirmed that deletion of *thiL* is lethal to *P. aeruginosa* unless TPP is exogenously provided in the media (Fig. 1). Complementation with the plasmid expressing *thiL* (p*thiL*) relieved the growth defect of the Δ*thiL* mutant caused by TPP depletion, suggesting that ThiL is essential for TPP biosynthesis in *P. aeruginosa* (Fig. 3A).

**Figure 1.**
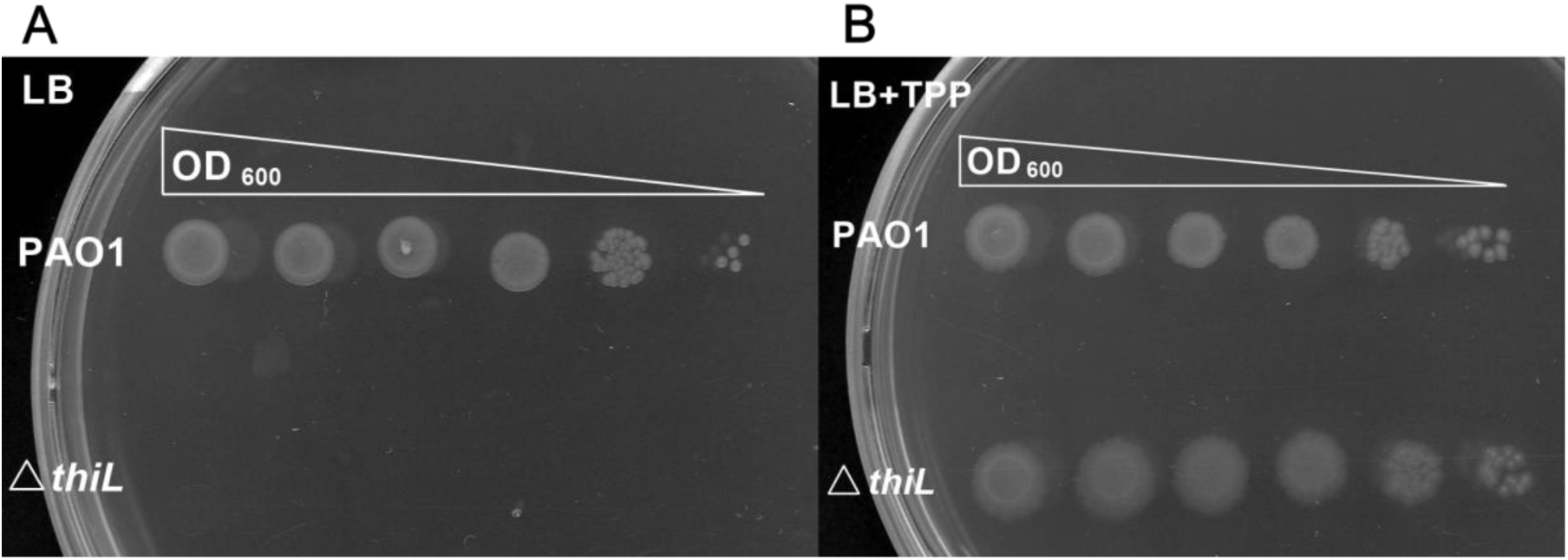
Phenotypic analysis of *thiL* deletion in *P. aeruginosa*. (A) A serial dilution of *thiL* deletion mutant (∆*thiL*) of *P. aeruginosa* was plated on LB agar, and showed no growth after 24-hours incubation at 37°C. (B) TPP supplementation (1mM) in LB agar rescued the growth defect of ∆*thiL*.

The thiamine salvage pathway is another way to generate TPP in many organisms, including bacteria and even mammals, which cannot synthesize TPP de novo. Two types of direct thiamine salvage pathways have been identified in bacteria: one-step pyrophosphorylation of thiamine to TPP by thiamine pyrophosphokinase (TPK/ThiN); and two-steps of subsequent monophosphorylation of thiamine to TMP then TPP by ThiK and ThiL, respectively. As the thiamine salvage pathway in *P. aeruginosa* has yet to be characterized and no thiamine transporter genes have been identified in this species (26), we first tested whether the bacteria is capable of producing TPP using extracellular thiamine. We found that the *thiE* mutant in which de novo TPP synthesis is impaired was able to grow in the presence of extracellular thiamine, TMP, and TPP, indicating that *P. aeruginosa* can take up thiamine derivatives and has a thiamine salvage pathway (Fig. 2). Unlike the *thiE* mutant, the Δ*thiL* strain exhibited a growth defect even with exogenously provided thiamine and TMP (Fig. 3). This result suggests that ThiL is involved in not only de novo TPP synthesis but also the thiamine salvage pathway in *P. aeruginosa*, indicating that ThiL plays a critical role in *P. aeruginosa* thiamine metabolism.

**Figure 2.**
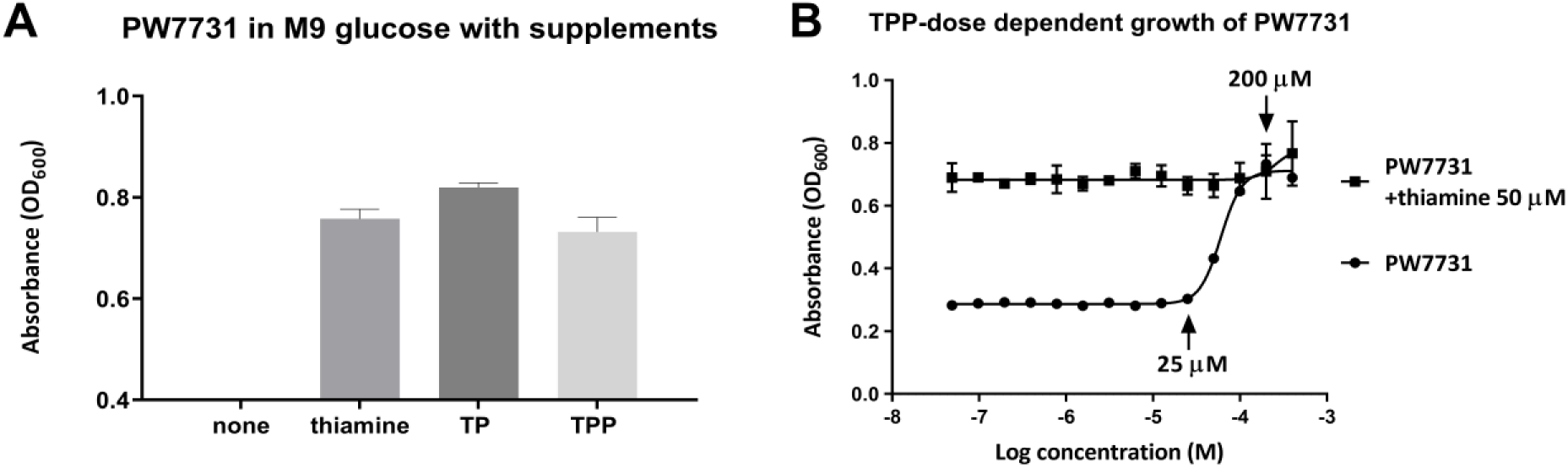
Growth of *thiE* mutant (PW7731) of *P. aeruginosa* in M9 glucose supplemented with thiamine, TMP, or TPP. (A)Absorbance (OD_600_) of PW7731showed normal growth in minimal media supplemented with thiamine (50µM), TMP (50µM), or TPP (100µM). (B)TPP dose-dependent growth of PW7731 revealed the salvage concentration range (25-200µM). Error bars are means ± standard deviation of triplicate experimental groups.

**Figure 3.**
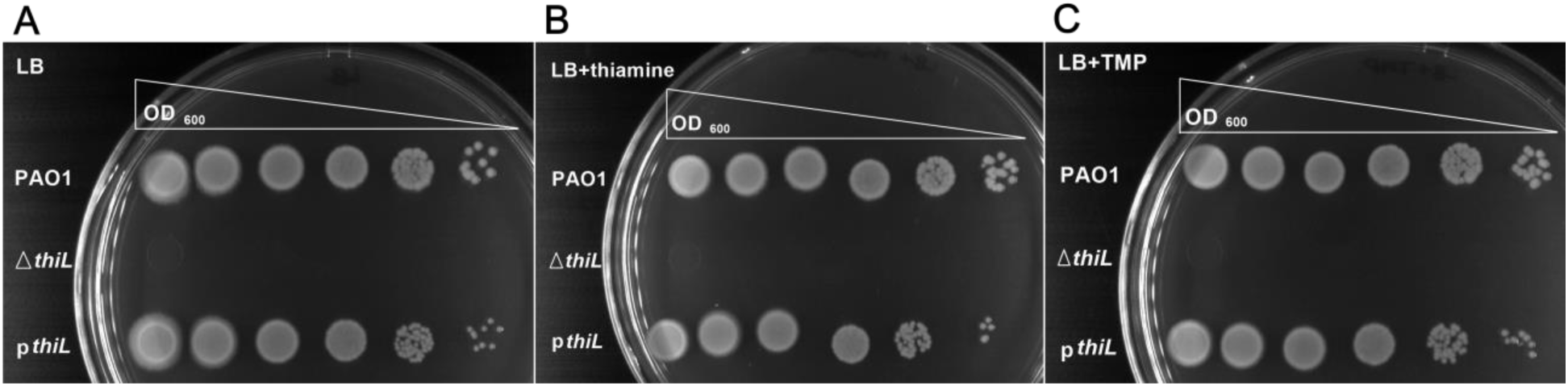
Phenotypic analysis of ∆*thiL* and *thiL* complementation by pBSPII SK(-)-*thiL* plasmid in *thiL* deletion mutant (p*thiL*) with thiamine or TMP supplementation. ∆*thiL* did not grow on (A) LB and LB supplemented with (B) thiamine (50 µM) or (C) TMP (50 µM) after 24 hours incubation at 37°C. p*thiL* showed normal growth in (A) LB and LB supplemented with (B) thiamine or (C) TMP.

### The role of ThiL in the virulence of P. aeruginosa in vivo

Next, we sought to determine the role of ThiL in the virulence of *P. aeruginosa* in vivo. C57BL/6 mice were infected intranasally with 2×10^7^ CFU of the PAO1, Δ*thiL*, or *thiL* complemented strains. During the first 20 hours, all infected mice exhibited decreased movement compared to uninfected mice. Histopathologic analysis of the lung tissues at 20-h post-infection also suggested that unlike uninfected control mice, all infected mice showed distinct inflammatory response composed mainly of peribronchial and alveolar neutrophilic infiltrates with neutrophilic consolidation and smooth muscle hyperplasia in the arterioles of the lung tissues (Fig. 4). Although mice infected with the Δ*thiL* strain exhibited a slightly lower degree of lung neutrophil infiltration (8.2±6.3%) than mice infected with the PAO1 strain (11±4%), the difference was not significant, suggesting that the Δ*thiL* mutant is capable of recruiting host immune cells and causing lung inflammation similar to the PAO1 strain (Fig. S2). Interestingly, when the average bacterial load in the left lobe of the lungs was measured at 20-h post-infection (n=5 for each group), over 1000 times more bacteria were found in mice infected with the PAO1 strain (1.6 × 10^8^ CFU) and *thiL* complemented strain (1.6 × 10^8^ CFU) compared with mice infected with the Δ*thiL* strain (6.7×10^4^ CFU) (Fig. 5a). In addition, the survival rates of the infected mice clearly demonstrated the attenuated virulence of the Δ*thiL* strain compared with the PAO1 and *thiL* complemented strains. Over 50% of mice infected with the PAO1 and *thiL* complemented strains died by 24-h post-infection, and the rest of the infected mice were dead by 48-h post-infection (Fig. 5b). In contrast, uninfected control mice and mice infected with the Δ*thiL* mutant survived as long as 72-h post-infection (Fig. 5b).

**Figure 4.**
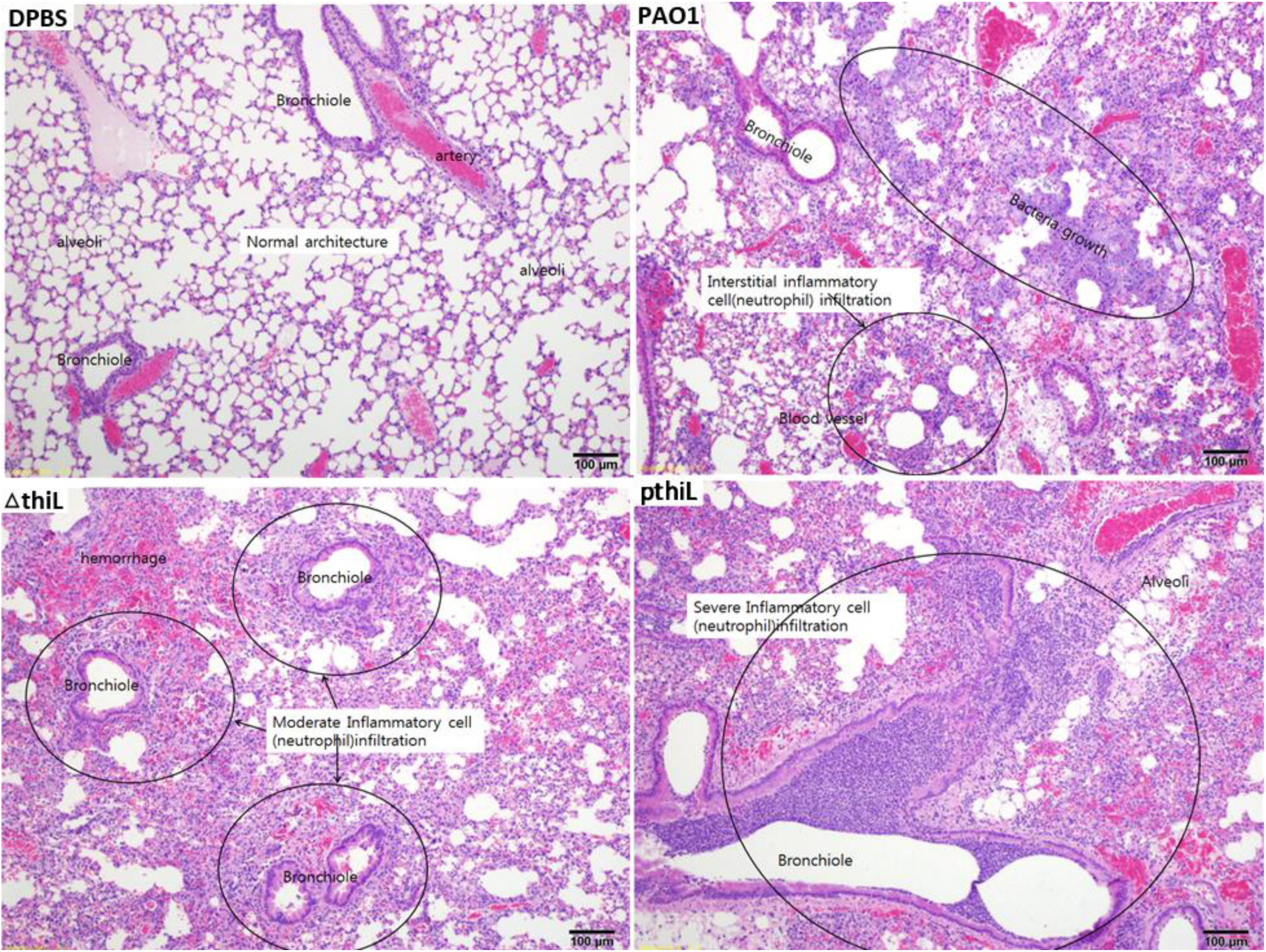
Neutrophile infiltration of the left lobe of lungs in C57BL/6 mice infected with PAO1, Δ*thiL*, or p*thiL*. Intranasal infections were applied to 6 weeks old mice with DPBS, PAO1, Δ*thiL*, or p*thiL*. H&E staining showed moderate to severe neutrophil infiltration (marked in circles) in bronchioles and blood vessels of the lungs after 20 hours of the infections. Magnification x100, Representative images of triplicate experimental groups.

**Figure 5.**
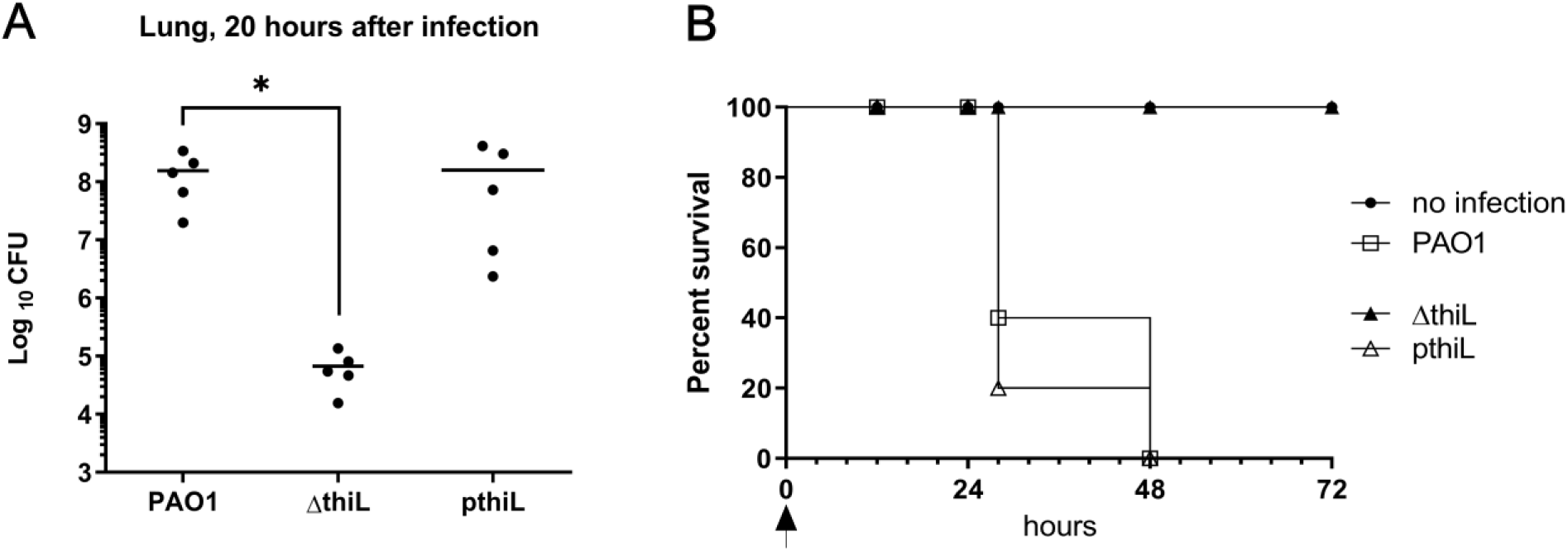
Bacterial loads in mice lungs infected with PAO1, Δ*thiL*, or p*thiL*, and survival analysis of the each infection groups. (A) CFU analysis showed relatively low bacterial loads in the lungs of mice infected with ∆*thiL* (6.7×10^4^) compared to PAO1 (1.6×10^6^), or p*thiL* (1.6×10^6^) after 20 hours of the infections. *, p<0.05 (B) 100% survival rate was observed in the mice infected with ∆*thiL* for 72 hours after the infections. The survival rates were reduced to below 50% in the mice infected with PAO1 or p*thiL* strains in 24 hours after the infection. Arrow(↑) indicated the time of infection. n=5 for each experimental groups, repeated three times.

As there was a discrepancy between the results of the lung histology and survival analyses, we further investigated pathogenesis in the infected mice by measuring the host immune response in the blood. Levels of three pro-inflammatory cytokines—IL-6, IL-8, TNF-α— were measured in the blood of infected mice at 20-h post-infection. The level of MIP-2 was measured as a murine counterpart of IL-8. Levels of all three tested cytokines were significantly higher in PAO1-infected mice than Δ*thiL*-infected mice (Fig. 6a,b,c). Similarly, the splenic bacterial count in mice infected with the PAO1 strain (6 × 10^5^ CFU/organ) was four orders of magnitude higher than that of Δ*thiL*-infected mice (2 × 10^1^ CFU/organ) (Fig. 6d). Taken together, these results suggest that overall virulence is attenuated in the Δ*thiL* strain due to decreased severity of bacteremia and sepsis that ultimately lead to the mice survival demonstrating that ThiL is essential for the full virulence of *P. aeruginosa*.

**Figure 6.**
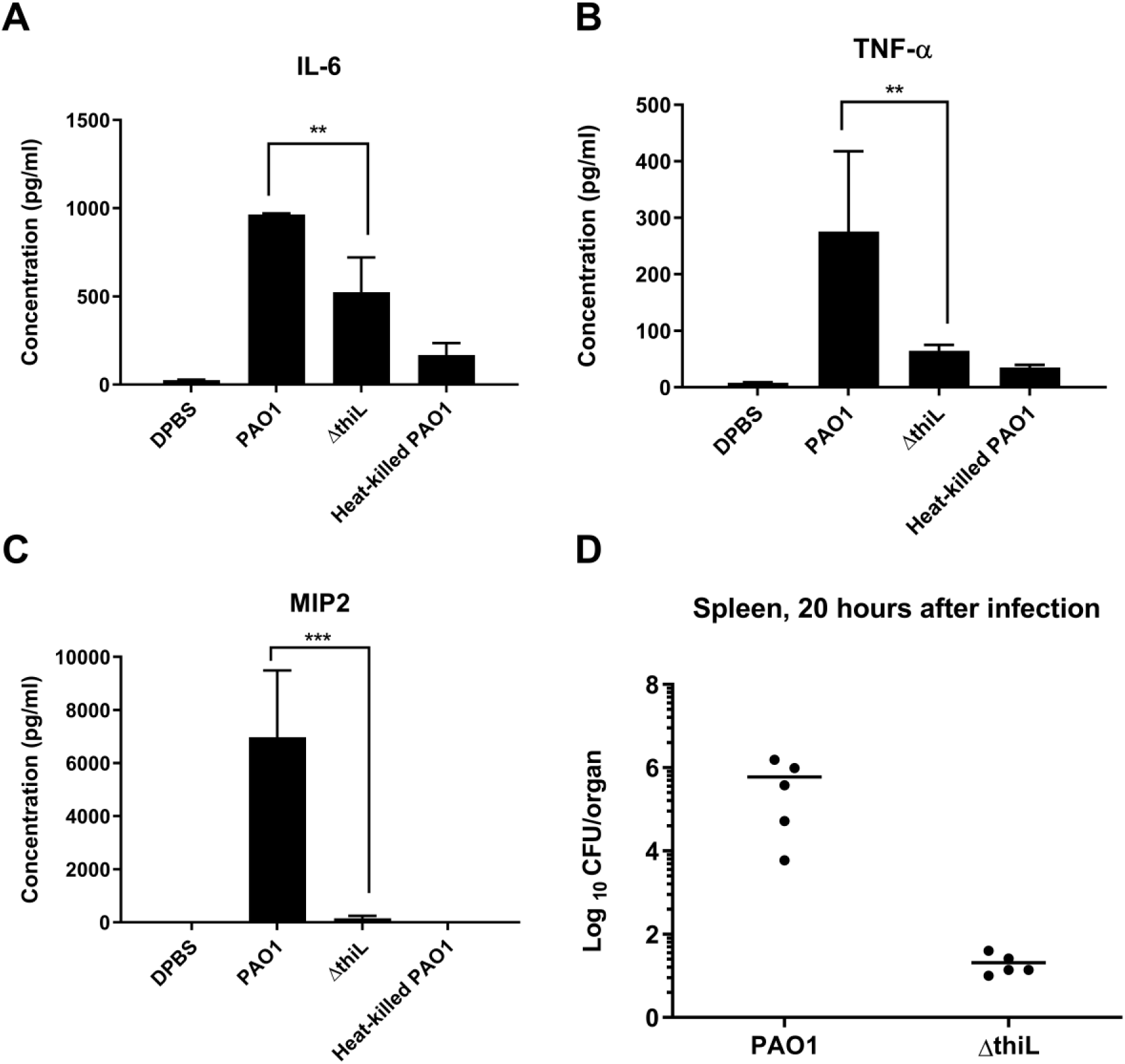
Pro-inflammatory cytokines (IL-6, TNF-α, MIP-2) of the mice blood after 20 hours of infection with PAO1, Δ*thiL*, or heat-killed PAO1, and bacterial loads (CFU/organ) in mice spleen after 20 hours of infection with PAO1 or Δ*thiL*. ELISA for (A) IL-6, (B) TNF-α, and (C) MIP-2 of mouse plasma indicated significantly low cytokine levels in Δ*thiL* groups compared to PAO1. **, p<0.01; ***, p<0.001 (D) CFU analysis showed 10^4^ time lower bacterial loads in spleen of the mice infected with Δ*thiL* compared to PAO1. n=5 for each experimental groups, repeated three times.

### Biochemical characterization of P. aeruginosa ThiL

Despite its importance in bacterial physiology and virulence, to our knowledge, *P. aeruginosa* ThiL has never been studied before, in neither molecular nor biochemical level. For biochemical characterization of the enzyme, the *P. aeruginosa thiL* gene (PA4051) was PCR amplified and cloned into pET-28b. The recombinant *P. aeruginosa* ThiL (PaThiL) was purified as an approximately 37-kDa His-tagged protein using Ni-NTA affinity matrix, with >95% purity (Fig. S3). We found that purified PaThiL was highly unstable without desalting, exhibiting 80% and 56% activity loss within 24 h at 4°C and −210°C, respectively (Fig. S4). A further gradual decrease in activity was observed over time. Therefore, purified PaThiL was frozen and stored in liquid nitrogen (−210°C) after immediate desalting, and activity was retained for over 1 month without significant loss (Fig. S4).

Previous studies of ThiL used HPLC-based assay or a coupled assay using apo-carboxylase to assess ThiL activity (28,36,40). Because both types of methods are quite inconvenient and inefficient, we adapted a luminescent kinase assay that detects ATP consumption and optimized it to determine PaThiL enzymatic kinetics (see the Materials & Methods section). The final assay was established to analyze the activity of 10 µg of PaThiL in reaction buffer containing 0.05 mM TMP, 0.05 mM ATP, 50 mM Tris-HCl (pH 8.0), 5 mM MgCl_2_, and 350 mM KCl. The K_M_ values for TMP and ATP, as well as the Vmax value, were calculated based on Michaelis-Menten and Lineweaver-Burk plots of PaThiL activity in reactions containing several different concentrations of ATP and TMP. The Vmax value of PaThiL was 4.0±0.2 nmol·min^−1^. The K_M_ values for ATP and TMP were 111±8 μM and 8.0±3.5 μM, respectively, in a random Bi-Bi mechanism (Fig. 7). Similar ranges of K_M_ values were previously reported for partially purified *E. coli* ThiL: 270 and 1.1 μM for ATP and TMP, respectively (36). Further characterization of PaThiL revealed that the enzyme phosphorylates oxythiamine monophosphate (a TMP analogue), with a K_M_ value of 15.2±2.0 μM (Fig. S5). Thiamine, as well as other thiamine analogues, including oxythiamine, pyrithiamine and amprolium, were not phosphorylated by PaThiL, indicating that prior acquisition of monophosphate is a requirement for PaThiL substrates (Fig. S6). TPP was not further phosphorylated by ThiL, suggesting that PaThiL does not contribute to the generation of intracellular thiamine triphosphate.

**Figure 7.**
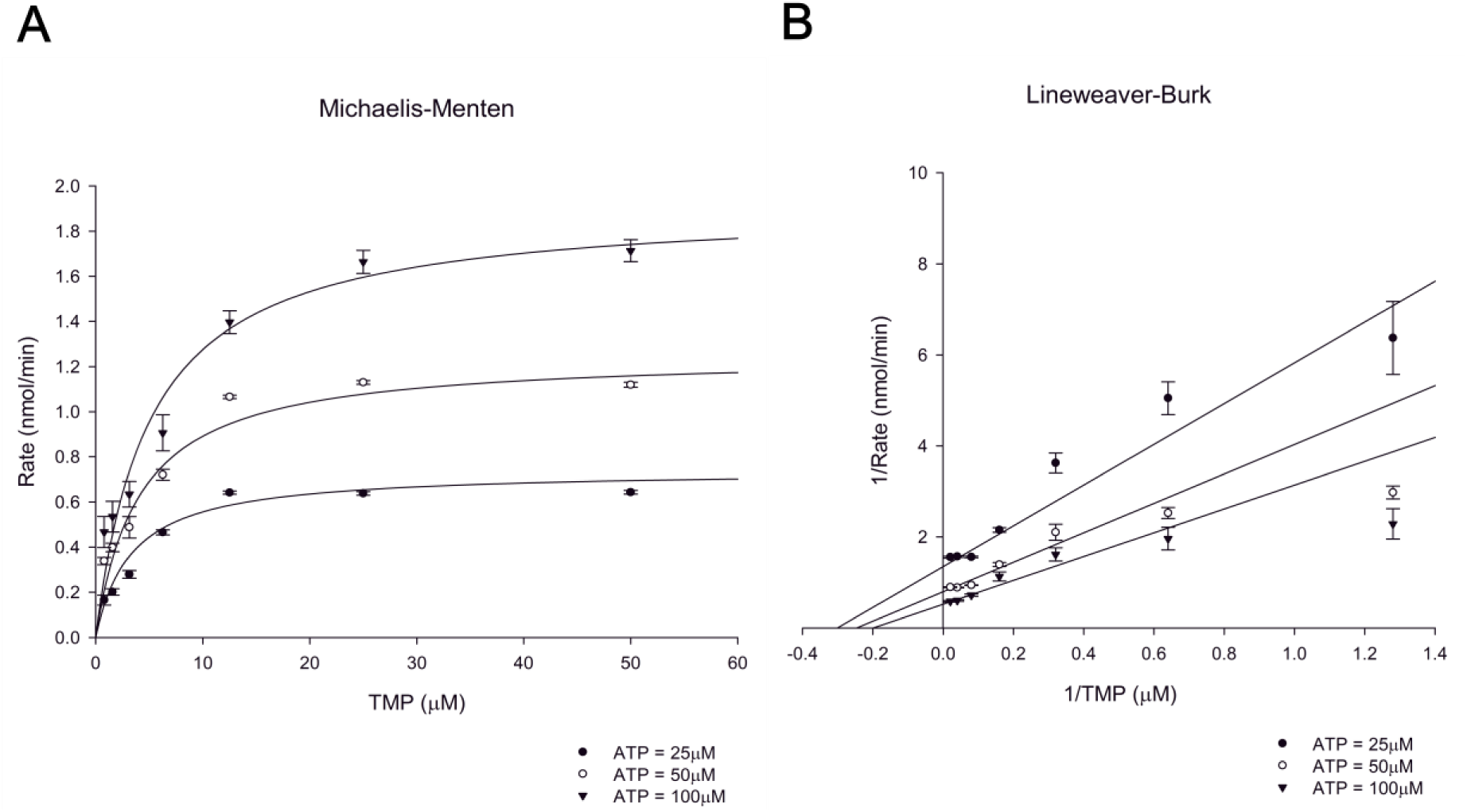
Kinetic parameter of ThiL by measuring the rate of kinase activity. Enzymatic assays were performed for isolated ThiL (10 µg) with various concentrations of TMP (0 to 50 µM). The assays were initiated with three ATP concentrations (25, 50, 100 µM) to determine ThiL activity rates. TMP data were plotted in (A)Michaelis-Menton and (B)Lineweaver-Burk functions. The result indicated K_M_ of ThiL for ATP (111±8), TMP (8.0±3.5), and Vmax (4.0.±0.2). Error bars are means ± standard deviation of triplicate experimental groups.

### Validation of PaThiL as a new antibacterial target

Gram-negative bacteria such as *P. aeruginosa* are notorious for their multi- and pan-drug resistance, which has created an urgent need for new classes of antibiotics. Our in vivo virulence study of the Δ*thiL* mutant indicated that ThiL is a promising novel candidate for antibacterial molecule, as humans have no PaThiL homolog. In order to validate druggability of PaThiL, approximately 2,800 commercially available compounds were screened for inhibitory activity against PaThiL. The screening was conducted using a slight modification of our established in vitro PaThiL assay. The screening assay was initiated upon addition of 5 µg purified PaThiL to a reaction mixture containing 10 µM TMP and 100 µM test compound; ThiL activity was measured by monitoring ATP consumption during a 10-min reaction. WAY213613 (Tocriscreen) and 5-hydroxyindolacetic acid (Lopac) were among several compounds that inhibited PaThiL. Further biochemical studies revealed that WAY213613 is a noncompetitive inhibitor of TMP for PaThiL, with a Ki value of 13.4±2.3 µM, whereas 5-hydroxyindolacetic acid is an uncompetitive inhibitor, which exhibited a Ki value of 114±27 µM (Fig. 8). Additional binding mode analyses of each compound with the homology modeling derived 3D structure of PaThiL suggested energetically favorable interactions with PaThiL (data to be published).

**Figure 8.**
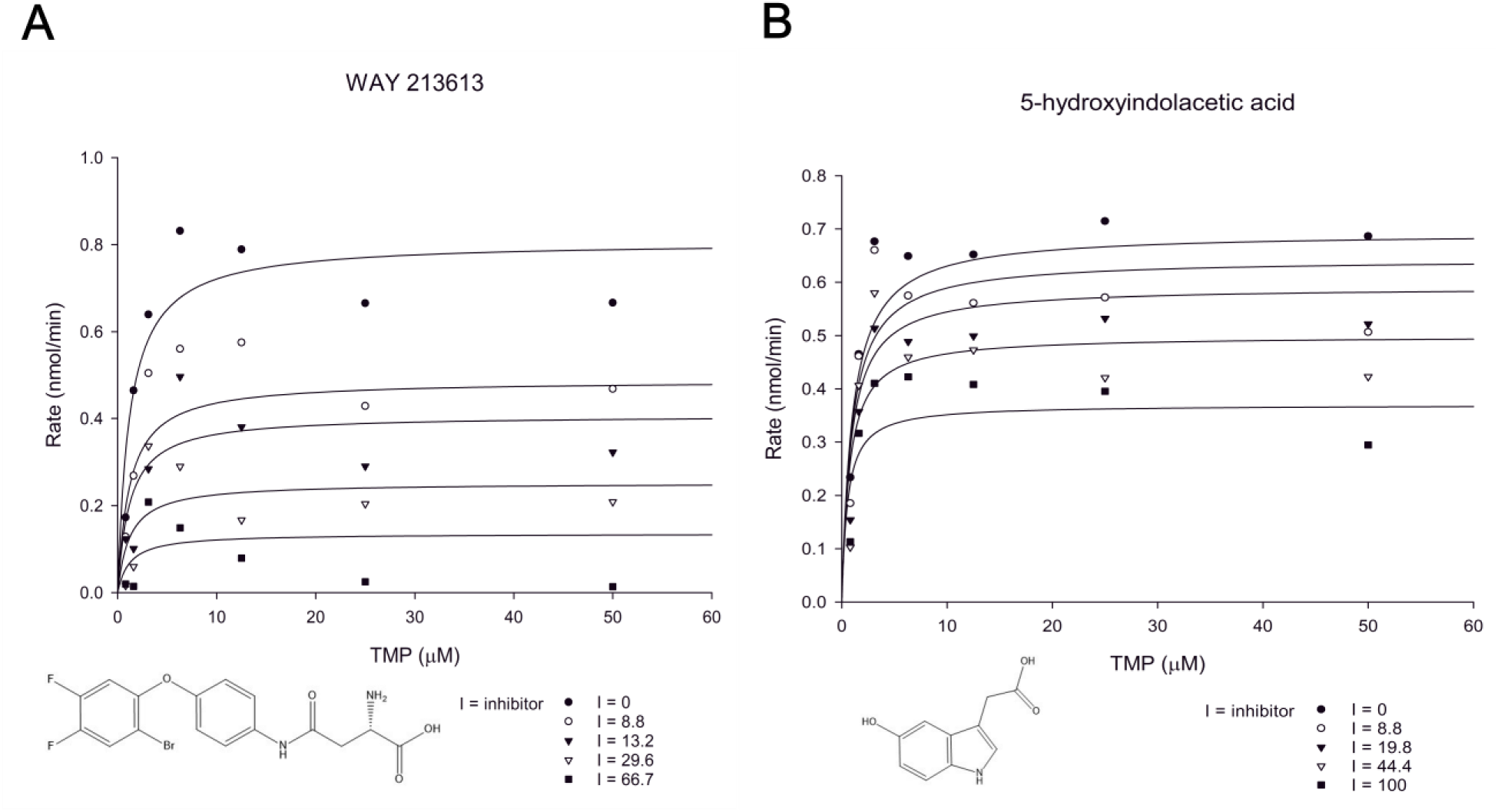
Enzymatic assay of ThiL with ThiL inhibitors for inhibition kinetics and Ki. Various concentrations of WAY213613 (0 to 66.7 µM) or 5-hydroxyindolacetic acid (0 to 100µM) was applied to ThiL assay with ATP (100 µM) and TMP (0 to 50 µM) to determine ThiL activity rates. The activity rates of ThiL with (A) WAY213613 or (B) 5-hydroxyindolacetic acid were plotted in Michaelis-Menton functions. The kinetic parameters showed that WAY213613 is a noncompetitive inhibitor with Ki value of 13.4±2.3 µM, and 5-hydroxyindolacetic acid is an uncompetitive inhibitor with Ki value of 114±27 µM.

We next tested whether the identified PaThiL inhibitors exhibit antibacterial activity. Because the outer membrane of *P. aeruginosa* can impede the entry of drugs into the cell and thereby mask potential antibacterial activity of PaThiL inhibitors, we also tested antibacterial activity with a low concentration of colistin (0.5 µg/ml), which can increase cell permeability by disrupting the bacterial outer membrane while not affecting bacterial viability (Fig. S7; Tab.S1). Although 5-hydroxyindoleacetic acid did not exhibit antibacterial activity at 100 µM even in the presence of colistin, we found that 100 µM WAY213613 completely ceased bacterial growth not by itself but in the presence of colistin (Fig. 9). These results suggest that PaThiL is a valid target for development of new antibacterial therapeutics demonstrating the feasibility of identification of small molecules that inhibit PaThiL.

**Figure 9.**
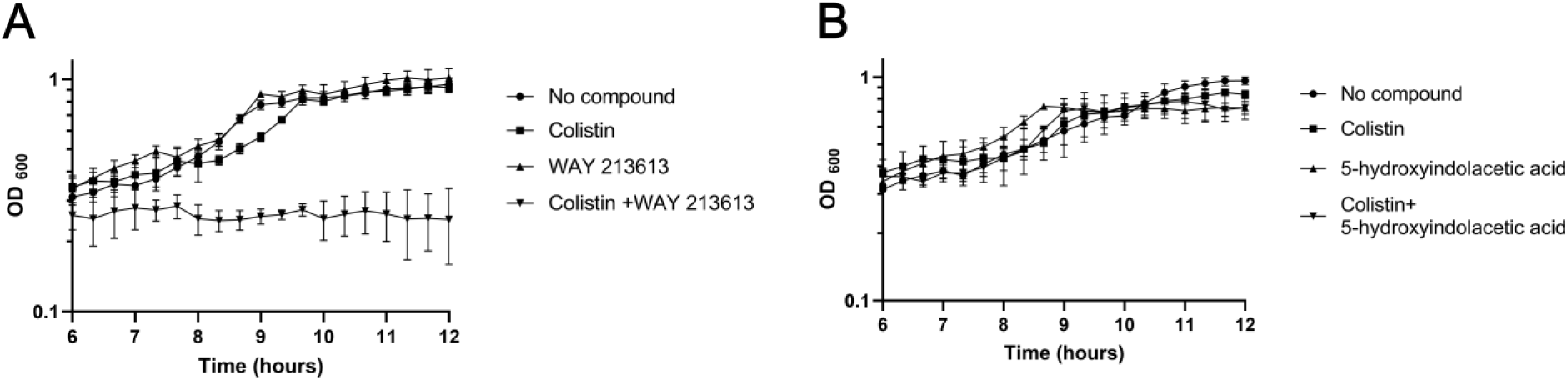
Growth kinetics of PAO1 for antibacterial effect test of WAY213613 or 5-hydroxyindolacetic acid. No antibacterial activities were observed with 100 µM of (A) WAY213613(▲) or (B) 5-hydroxyindolacetic acid (▲). Bacterial inhibition effect for (a)WAY213613 (100 µM) was observed with addition of a sublethal dosage of colistin (0.5 µg/ml) (▼). (b)5-hydroxyindolacetic acid (100 µM) did not show antibacterial effect with addition of colistin (0.5 µg/ml)(▼). Control groups were no compound in LB (●) or LB with colistin 0.5 µg/ml (■). Error bars are means ± standard deviation of triplicate experimental groups

## Discussion

TPP-dependent enzymes including transketolase, pyruvate dehydrogenase, 2-oxoglutarate dehydrogenase, and 1-deoxy-d-xylulose-5-phosphate synthase, catalyze essential cellular reactions in bacteria, from central metabolism to biosynthesis of amino acids, cofactors, and lipids (21). Recently, TPP-dependent pyruvate and α–ketoglutarate dehydrogenase complexes in *P. aeruginosa* were found to intracellularly reduce phenazin derivatives such as pyocyanin (a pseudomonal toxin), thus contributing to iron acquisition and redox homeostasis of the bacteria in addition to their designated roles in central metabolism (41). This evidence indicates that TPP deficiency can have a substantial effect on the metabolic network of bacteria, thus making TPP metabolism a very attractive target for the development of new antibiotics. Indeed, inhibitors of ThiE exhibit antibacterial activity against *Mycobacterium tuberculosis*, a notorious human pathogen (42). As *M. tuberculosis* does not harbor the genes for the thiamine salvage pathway and transporters, inhibition of ThiE can effectively deplete the TPP pool by impairing TPP biosynthesis, the only source of TPP in the bacteria, ultimately causing the bacterial death.

However, inducing the death of many pathogenic bacteria via TPP deficiency is hampered by several challenges, particularly in vivo: 1) complete deficiency of TPP requires concurrent inhibition of both thiamine salvage and TPP biosynthesis, and 2) levels of thiamine derivatives in the host, specifically TPP, should not be sufficient for directly promoting bacterial survival. In this work, we addressed these two challenges by investigating ThiL in *P. aeruginosa*. We demonstrated that ThiL is the key enzyme for both de novo TPP synthesis and thiamine salvage in *P. aeruginosa* by revealing an in vitro growth defect of the *thiL* mutant, thus indicating a possibility to deplete the TPP pool in *P. aeruginosa* by targeting ThiL (Fig. 1).

Although thiamine transporter genes have not been identified in *P. aeruginosa*, the growth of *thiE* and *thiL* mutants in the presence of thiamine, TMP, or TPP suggests the existence of transporters that import thiamine derivatives in *P. aeruginosa* (26) (Figs. 2–3). In *E. coli*, intracellular TPP concentrations were measured at 150~300 µM under optimal growth conditions, which are presumably TPP levels for normal bacterial growth (43,44). Consistent with this notion, when we determined the extracellular TPP concentration necessary to relieve the growth defect of the *thiE* mutant in which the TPP biosynthetic pathway is impaired, we found that a minimum TPP concentration of 25 µM was required to detect any degree of *thiE-*mutant growth, and a concentration over 200 µM was required to fully support the growth of the *thiE* mutant (Fig. 2). This result suggests that if *P. aeruginosa* relies solely on exogenous TPP for growth, the concentration should be higher than 25 µM. Reported TPP levels in mouse blood range from 0.3 to 1.2 µM, well below the TPP concentration sufficient to support growth of the *thiL* mutant (45,46). Consistent with these data, we found that the *thiL* mutant exhibited markedly reduced pathogenicity in a mouse infection model when compared with the wild-type strain (Figs. 5–6). TPP levels in healthy human blood (116~138 nM) are even lower than in mouse blood, suggesting that ThiL is a viable therapeutic target in the treatment of *Pseudomonas* infections (47,48).

We also validated *P. aeruginosa* ThiL (PaThiL) as a suitable target for new antibacterial agents by conducting an intensive biochemical characterization followed by identification of inhibitors of PaThiL. Considering the high intracellular concentrations of ThiL substrates (ATP: ~1.8 mM, TMP: ~12.5 µM) and product (TPP: ~0.3 mM) in bacteria (44), noncompetitive ThiL inhibitors might be more beneficial than competitive inhibitors that probably require a very high affinity for ThiL. Interestingly, we found that WAY213613, a known inhibitor of EAAT2 glutamate transporter, can inhibit PaThiL in a noncompetitive manner, with a Ki value of 13.4±2.3 µM. When we tested antibacterial activity, WAY213613 was able to inhibit the growth of *P. aeruginosa*, although the permeability of the bacterial outer membrane had to be increased to detect the activity (Fig. 9).

Antibiotic-resistant bacteria such as pan-drug resistant *P. aeruginosa* pose an immediate threat to global public health and highlight an urgent need for new antibiotics and targets. In this work, we extensively characterized ThiL of *P. aeruginosa* and demonstrated its roles in bacterial physiology and pathogenesis. Although this study focused on *P. aeruginosa* ThiL, this enzyme is expected to play similar roles in other bacteria that have a thiamine salvage pathway composed of ThiK and ThiL, such as *E. coli* and *Salmonella* species (26). Taken together, the results of this work demonstrate that ThiL is a suitable therapeutic target for the development of new drugs to treat not only *P. aeruginosa* infections but potentially many other bacterial infections as well.

## Experiment procedures

### Bacterial strains, culture media, and chemicals

All strains and plasmids used in this study are listed in Table 1. Bacteria were cultured in Luria-Bertani (LB) broth at 37°C with agitation. *thiE* transposon mutant (PW7731) was cultured in minimal media supplemented with thiamine monophosphate (TMP) for normal growth. LB powder was dissolved in purified water and autoclaved at 121°C for 15 minutes. As minimal media, M9 salt powder was dissolved in purified water and autoclaved at 121°C for 15 minutes. Sterile MgSO_4_ (2mM) and CaCl_2_ (0.1mM) were added to M9 solution for complete minimal media. Supplementations of the media including glucose (20mM), thiamine, TMP and TPP were sterilized by filtration through 0.22-µm PVDF membrane (Millipore, Burlington, MA, USA) before addition. Agar (1.5%) was added the media to prepare solid growth plates. Kanamycin (50 mg/ml) and ampicillin (100 mg/ml) were dissolved in purified water and sterilized by filtration through 0.22-µm PVDF membrane. Antibiotic stock solutions were stored at −20°C. Isopropyl β-D-1-thiogalactopyranoside (IPTG) (1M) was dissolved in purified water and filtered through a 0.22-µm filter. All chemicals were purchased from Sigma-Aldrich (St. Louis, MO, USA), except for ATP and IPTG, which were purchased from Thermo Fisher (Waltham, MA, USA).

**Table 1.**
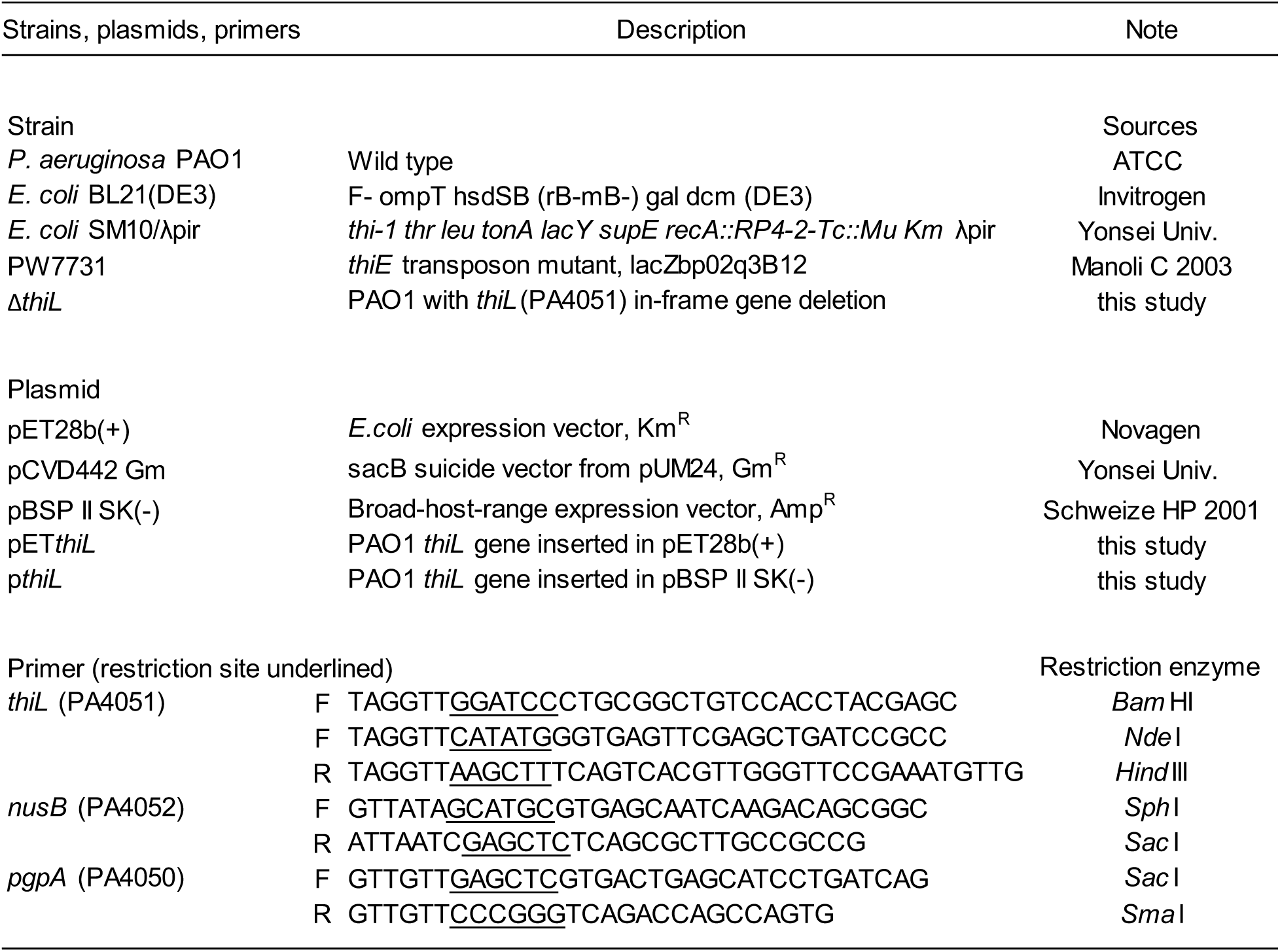
Strains, plasmids, primers used in this study

### Growth of thiE transposon mutant (PW7731) with thiamine, TMP, or TPP

M9 glucose was supplemented with thiamine (50 µM), TMP (50 µM), or TPP (100 µM), and the media were applied to PW7731 with initial OD_600_ of 0.02. After 16-hour incubation with agitation, absorbance (OD_600_) was measured for each growth condition. TPP salvage concentration was determined by 2-fold serial dilution of TPP in a range from 0.02 to 400 µM. Each concentration of TPP was applied to PW7731 with initial OD_600_ of 0.0001, which was prepared by 1/500 dilution of the bacteria at OD_600_ of 0.05 and incubated in 37°C for 16 hours. The absorbance (OD_600_) was measured with a multi-label plate reader.

### Overexpression and isolation of ThiL

The gene encoding thiamine monophosphate kinase (*thiL*) was amplified by PCR analysis using the primers listed in Table 1 and inserted into the pET28b(+) (Novagen, Madison, WI, USA), a vector for the cloning and expression of recombinant proteins with His-tags, at restriction sites between *Nde*I-*Hin*dIII. The plasmid was transformed into *E. coli* BL21 cells, which were selected on media containing kanamycin (50 µg/ml). To induce protein expression, IPTG (1 mM) was added to the mid-log phase culture and incubated for 2.5 hours at 37°C with agitation. The cells were centrifuged at 4000 × *g*, and the resulting pellet was resuspended in 5 ml Tris-HCl (100 mM [pH 8.0]) supplemented with a protease inhibitor (Roche, Mannheim, Germany). The suspended cells were then sonicated using a Bioruptor sonication system (Diagenode, Denville, NJ, USA) and centrifuged at 14,000 × *g*. ThiL protein in the supernatant was purified by affinity chromatography (GE Healthcare, Freiburg, Germany). The collected protein was desalted using a 30-K centrifugal filter (Millipore, Burlington, MA, U.S.A) and suspended in Tris-HCl (100 mM [pH 8.0]). The amount of protein in the collection was quantified using microassay protocol of Bradford protein assay with bovine serum albumin (BSA) as a standard. The protein was aliquoted in cryotubes and stored in −210°C for long-term preservation. The purity of the isolated protein was over 95% which was confirmed by Coomassie Blue staining.

### ThiL assay

The ThiL assay was established based on previous protocols (28,40). The assay reaction buffer contained 0.05 mM TMP, 0.05 mM ATP, 50 mM Tris-HCl (pH 8.0), 5 mM MgCl_2_, and 350 mM KCl. A total of 10 µg ThiL protein was added to the reaction buffer to initiate the kinase reaction. After 10 minutes, the amount of ATP was determined using a luminescence assay (Promega, Madison, WI, USA). ATP hydrolysis in the reaction buffer without TMP was less than 1% in 10 minutes. K_M_ and Vmax values were determined for TMP and ATP using the ThiL assay with TMP and ATP ranging from 0.78 to 50 and 25 to 100 µM in two-fold dilution steps, respectively. The assay data were fit to Michaelis-Menton or Lineweaver-Burk functions using the random bi-substrate module in Sigma Plot ver. 14.(Systat Software Inc. San Jose, CA, USA) to obtain the K_M_ and Vmax values. To determine the Ki of ThiL inhibitors, ThiL assays were carried out with WAY213613 (0 to 66.7µM) or 5-hydroxyindolacetic acid (0 to 100µM). The activity rates of ThiL were fit to Michaelis-Menton or Linewaver-Burk functions using single substrate-single inhibitor module in Simga Plot.

### Bacterial growth kinetics with ThiL inhibitors

Growth kinetics of PAO1 with ThiL inhibitors 100 µM of WAY213613 or 5-hydroxyindolacetic acid was applied to PAO1 with or without colistin 0.5 µg/ml. The growth kinetic assays were carried out at 37°C in 96-well plate with initial OD_600_ of 0.0001. Absorbance values (OD_600_) were measured every 20 minutes for 13 hours with a multi-label plate reader Spectramax M5 (Molecular Devices, San Jose, CA, USA). Checkerboard assay was also performed for combination of WAY213613 and colistin in the ranges of 0 to 200 uM and 0 to 20 uM, respectively. The MIC of each antimicrobial agent alone and in combination was defined as the lowest concentration that inhibited visible growth of PAO1. The fractional inhibitory concentration (FIC) index was calculated by the formula FIC= (MIC of WAY213613 in combination/MIC of WAY213613 alone) + (MIC of colistin in combination/MIC of colistin alone) as previously described (49,50).

### Construction of thiL deletion mutants via allelic exchange

To construct a *P. aeruginosa* mutant lacking *thiL*, allelic exchange was carried out according to previous studies (37,39). Briefly, both upstream (*nusB*, 480 bp) and downstream (*pgpA*, 516 bp) genes flanking *thiL* were amplified by PCR analysis using the primers listed in Table 1. Both ends of the amplified genes carried specific restriction sites (Table 1). The downstream gene and pCVD442 Gm, a suicide vector, were double digested with *Sac*I and *Sma*I and then ligated. Consecutively, the upstream gene and pCVD442-Gm-pgpA were double digested with *Sph*I and *Sma*I and then ligated to obtain the suicide vector carrying the amplified genes. The generated vector was electroporated into *E. coli* SM10 lambda *pir* to facilitate conjugation into *P. aeruginosa*. Transconjugants were selected on medium containing gentamicin (100 µg/ml), and allele exchange was induced by incubation in medium containing sucrose (6%). The deletion mutant was selected on LB medium supplemented with 1 mM TPP, and deletion of *thiL* was confirmed by PCR analysis.

### Generation of the complemented strain of the thiL deletion mutant

To generate a complemented strain of the *P. aeruginosa ΔthiL* mutant, the amplified *thiL* gene was double digested with *Bam*HI and *Hin*dIII and inserted into pBSP II SK(−) to create p*thiL*. The constructed plasmid was electroporated into competent cells of the PAO1 Δ*thiL* strain using a previously reported method (51). Briefly, the bacterial culture was washed twice and suspended with room temperature 300 mM sucrose to generate electrocompetent cells. Next, 1 µg p*thiL* was mixed with 100 µl electrocompetent cells, and the pulse was applied to the mixture (*P. aeruginosa* 2.5-kV setting, Bio-Rad) LB medium was then immediately added to the mixture and incubated for 1 hour at 37°C. Transformed cell were selected on LB agar plates supplemented with ampicillin (0.5 mg/ml). Complementation was confirmed by PCR analysis and bacterial growth without TPP supplementation.

### P. aeruginosa acute murine infection model

Inbred 6-week-old female C57BL/6 mice were purchased from Orient Bio (Seongnam, Korea) and maintained in our animal facility which cared the animals in accordance with the institutional guidelines.. All mice were subjected to a 1-week adjustment period prior to the experiment. The protocol was approved by the Animal Care and Ethics Committee of Institut Pasteur Korea (IPK-17014)

The acute infection model was established based on *Pseudomonas Methods and Protocols* (52). Bacteria were grown to mid-log phase and washed three times with Dulbecco’s phosphate-buffered saline (DPBS). Mice were anesthetized with a mixture of ketamine and rompun and inoculated intranasally with 2 × 10^7^ CFU bacteria suspended in 20 µl DPBS. At 20 hours after inoculation, mice were sacrificed using isofluorane, and both lungs and the spleen were harvested. The left lung and spleen were homogenized in PBS and then plated separately onto LB agar plates for wild-type and complementation strains, and the LB agar was supplemented with TPP for the Δ*thiL* strain. The plates were incubated for 20 hours at 37°C, and then the viable bacterial cells were counted to determine the number of bacteria in the organs. The cranial lobe of the lungs was placed in 10% formalin for histopathologic analysis by hematoxylin and eosin (H&E) staining. For histopathological analysis, 4 µm-thick sections of fixed lungs were blindly analyzed for cellular infiltration using Image-Pro® Plus ver 4.5, Media Cyberbetics (Rockvile, MD, USA). Briefly, degrees of neutrophil infiltration were assessed by percentages of inflammatory lesions per total areas in microscopic images (40x) of the lung sections. Another set of mice (n=5) was subjected to body weight measurement and clinical signs of infection, and significances were recorded until 72 hours after inoculation. At the end of the experiment, the mice were euthanized using isofluorane.

### Determination of pro-inflammatory cytokine plasma concentrations

At the time of sacrifice, an average of 0.7 ml whole blood was withdrawn from each mouse by cardiac puncture and collected in heparin-treated tubes. The blood was immediately centrifuged at 2000 × *g* for 20 minutes to obtain plasma. The isolated plasma was fast-frozen on dry-ice and stored at −80°C. To measure cytokine concentrations, we used Quantikine ELISA kits for interleukin 6 (IL-6), tumor necrosis factor– alpha (TNF-α), macrophage inflammatory protein 2 (MIP-2), and IL-1 beta (R & D Systems, Minneapolis, MN). Each assay was performed according to the manufacturer’s instructions.

### Data Availability

All data is contained within the article and in the online *Supporting Information*.

## Acknowledgements

We thank Prof. Sangsun Yoon for invaluable advice on this research. This work was supported by the National Research Foundation of Korea (NRF) grant funded by the Korea government (MSIT)(NRF-2017M3A9G6068246) as well as Gyeonggi-do.

